# Machine learning-assisted neurotoxicity prediction in human midbrain organoids

**DOI:** 10.1101/774240

**Authors:** Anna S. Monzel, Kathrin Hemmer, Tony Kaoma Mukendi, Philippe Lucarelli, Isabel Rosety, Alise Zagare, Silvia Bolognin, Paul Antony, Sarah L. Nickels, Rejko Krueger, Francisco Azuaje, Jens C. Schwamborn

## Abstract

A major challenge in the field of neurodegenerative diseases is the poor translation of pre-clinical models to clinical applications. The human brain is an immensely complex structure, which makes it difficult to recapitulate its development, function and disorders. In the recent years, brain organoids derived from human induced pluripotent stem cells have risen as novel tools to study neurodegenerative diseases such as Parkinson’s disease (PD). PD is a multifactorial disorder, with aging, genetics and environmental factors as key etiological elements. The majority of the PD cases are idiopathic and proposed to result from a complex interaction between genetic predisposition and environmental exposure. Consequently, the identification of potentially disease causing environmental factors is of critical importance. Organoids, as complex multi-cellular tissue proxies, are an ideal tool to study cellular response to environmental changes. However, with increasing complexity of the system, usage of quantitative tools becomes challenging. This led us to develop an automated high-content image analysis pipeline for image-based cell profiling in the organoid system. Here, we introduce a midbrain organoid system that recapitulates features of neurotoxin-induced PD, representing a platform for machine-learning-assisted prediction of neurotoxicity in high-content imaging data. This model is a valuable tool for advanced *in vitro* PD modeling and for the screening of putative neurotoxic compounds.

## Introduction

Parkinson’s disease (PD) is the second most-common neurodegenerative disorder with an estimated global prevalence of ∼6 million people that is expected to double by the year 2040 (Dorsey and Bloem, 2018; GBD, 2017). The disease is clinically diagnosed after the onset of primary motor symptoms including resting tremor, bradykinesia, rigidity and postural instability, when most of the dopaminergic neurons are degenerated. The subsequent depletion of dopamine in the striatum is the fundamental mechanism causing the motor features in PD. The etiology of PD is multifactorial, with endogenous (genetic) and exogenous (environmental) contributors to development and onset. While a minority (~10%) of the cases can be explained by well-defined genetic causes (Klein and Westenberger, 2012), in the majority of PD the cause is unknown. Accumulating evidence suggests that the etiology is only partially explained by the patient’s genetic background. It is proposed that the combination and interaction of hundreds of genetic risk variants, ageing, and environment leads to development and onset of PD, suggesting that "multiple hits" are necessary for the disease to develop (Bellou et al., 2016; Schwamborn, 2018; Sulzer, 2007). This highlights the necessity to expand the research to identify potential neurotoxic compounds and their harmful effects on the human brain.

One of the major challenges for PD is the development of reliable disease models that can capture the complex nature of the human brain and related disorders. Animal models with toxin-induced neurodegeneration or genetically modified organisms are still the gold standard in brain research. However, rodents cannot reproduce the complexity of the human brain because brain development, anatomy and physiology differs greatly between animals and humans (Hodge et al., 2019). Hence, findings are not always transferable to the human condition and the success rate of preclinical trials is very low. In addition, *in vivo* animal toxicity testing is constrained by ethical considerations, time and financial burdens. On the other hand classical two-dimensional *in vitro* cell culture models with isolated cell types are too uniform and homogeneous to model a complex organ such as the human brain. To bridge the gap between classical 2D *in vitro* and complex *in vivo* models, stem cell-derived 3D cell culture systems such as brain organoids have risen in the recent years. These complex *in vitro* systems effectively mimic the organ architecture and function, and have been shown to model neurodevelopmental and neurodegenerative disorders (reviewed in (Wang, 2018)). Notably, these models can be derived from human induced pluripotent stem cells, which makes them an ideal platform for personalized medicine.

However, with increasing complexity of the system, the availability of tools is limited due to enormous processing of the data. So far powerful techniques such as high-throughput screening have a limited relevance in the organoid system, because generating organoids is highly laborious and requires manual handling. Due to architecturally complex heterotypic organization, comprehensive image analysis in organoids is challenging. Building on this, we developed methods to automatically acquire and process data from high-content imaging in organoids, which has been successfully demonstrated in brain organoids (Smits et al., 2019) and 3D microfluidic cultures (Bolognin et al., 2019). In this study, we further refined this pipeline with optimized high-content image data analysis tools in a neurotoxin-induced PD organoid model. Furthermore, we used machine learning (ML) tools to complement the *in vitro* toxicity assay. ML is gaining popularity in toxicity prediction because computational methods can combine a variety of different measurements and information sources to predict an outcome of interest (Scheeder et al., 2018). In high content imaging data, the various sources can originate from cell type abundance, cellular morphology or degenerative features and cell death. With high-content imaging tools, we obtain high-resolution data on the single cell level and assessed the neurotoxic effect of the catecholaminergic neurotoxin 6-OHDA on human midbrain organoids using random forest classification. This pipeline, from treatment to prediction, is valuable and useful for the exploration of potential neurotoxic compounds in complex human brain organoids.

## Results

### Generation and 6-OHDA-treatment of midbrain organoids from hNESCs

To generate human midbrain organoids, we used a previously published method (Monzel et al., 2017) starting from human neuroepithelial stem cells (hNESCs). hNESCs were derived from human induced pluripotent stem cells (hiPSCs) from three healthy individuals (Table S1). To specifically target the dopaminergic system, organoids were exposed to the catecholaminergic neurotoxin 6-hydroxydopamine (6-OHDA) (Fig. 1). In dopaminergic neurons, 6-OHDA, due to its structural similarity with endogenous dopamine, exerts its toxic effects by crossing the dopamine transporter, leading to an accumulation of the toxin in the neuron (Emborg, 2007; Jonsson and Sachs, 1975; Thoenen and Tranzer, 1968; Ungerstedt, 1968).

**Fig. 1:**
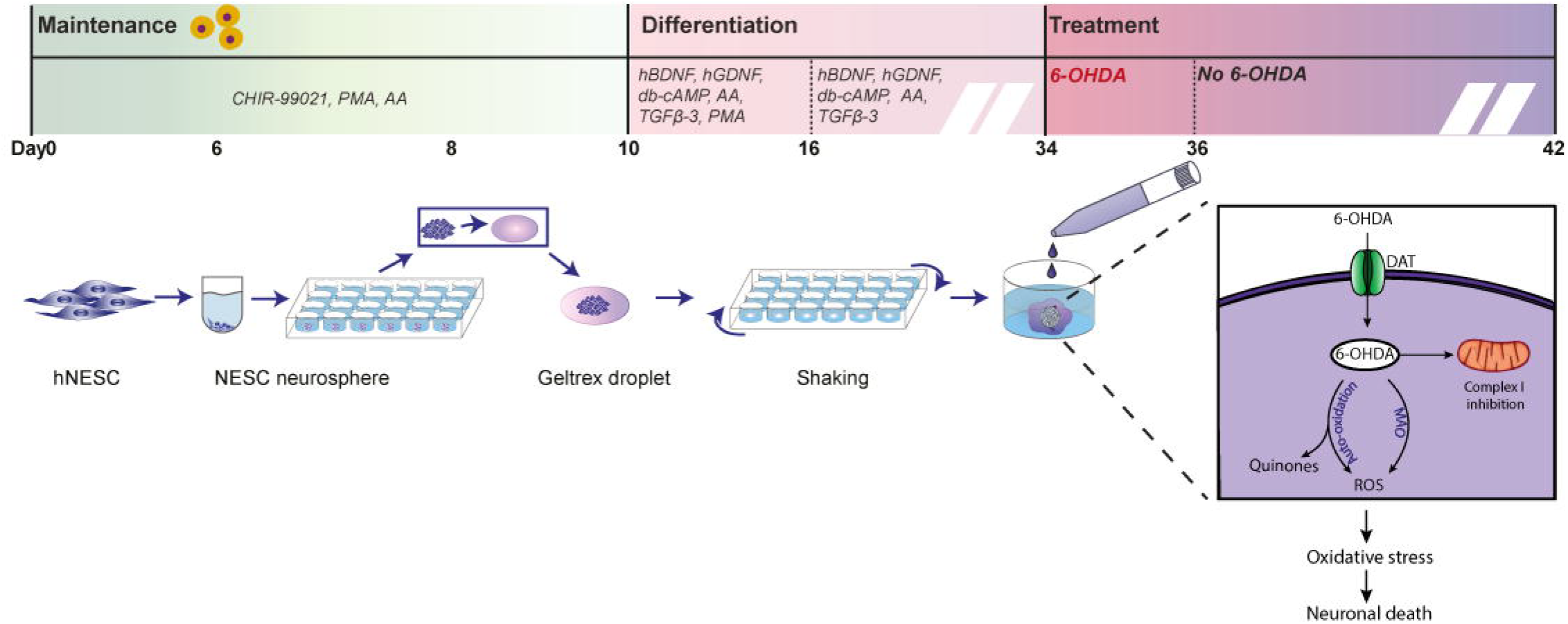
Experimental setup and mechanism of action of 6-OHDA. Human iPSC-derived neuroepithelial stem cells (hNESCs) were cultured under 3D conditions, embedded in droplets of Geltrex and differentiated into midbrain dopaminergic neurons after 10 days. Organoids were cultured on an orbital shaker (as described in (Monzel et al., 2017)). After 5 weeks of organoid culture, organoids were treated with different concentrations of 6-OHDA, leading to the formation of cytotoxic species, which is followed by neuronal death. (Adapted from (Monzel et al., 2017))

### 6-OHDA induces concentration-dependent cell death in midbrain organoids

We investigated 6-OHDA-induced neurotoxicity by treating organoids with various concentrations of 6-OHDA, ranging from 50µM to 500µM for 48h, followed by one additional week under normal culture conditions. Cell quantification by flow cytometry revealed that exposure to the toxin caused a concentration-dependent reduction in the amount of living dopaminergic neurons, identified by the rate-limiting enzyme of the dopamine synthesis, Tyrosine hydroxylase (TH) (Fig. 2a, b, Fig. S1a). We fitted a non-linear regression curve for each cell line and determined a mean LD_50_ at 147µM 6-OHDA (Fig. 2c). Since LDs varied among the cell lines, we used a concentration of 175µM in further experiments, which led to a significant reduction in the amount of dopaminergic neurons in all three cell lines (Fig. 2d). Consistent with the FACS data, we observed an overall concentration-dependent reduction in the TH protein in immunofluorescence staining (Fig. S1b) and Western Blot, resulting in an average 2.3 fold decrease of the protein after 175µM 6-OHDA treatment (Fig. 2e, f).

**Fig. 2:**
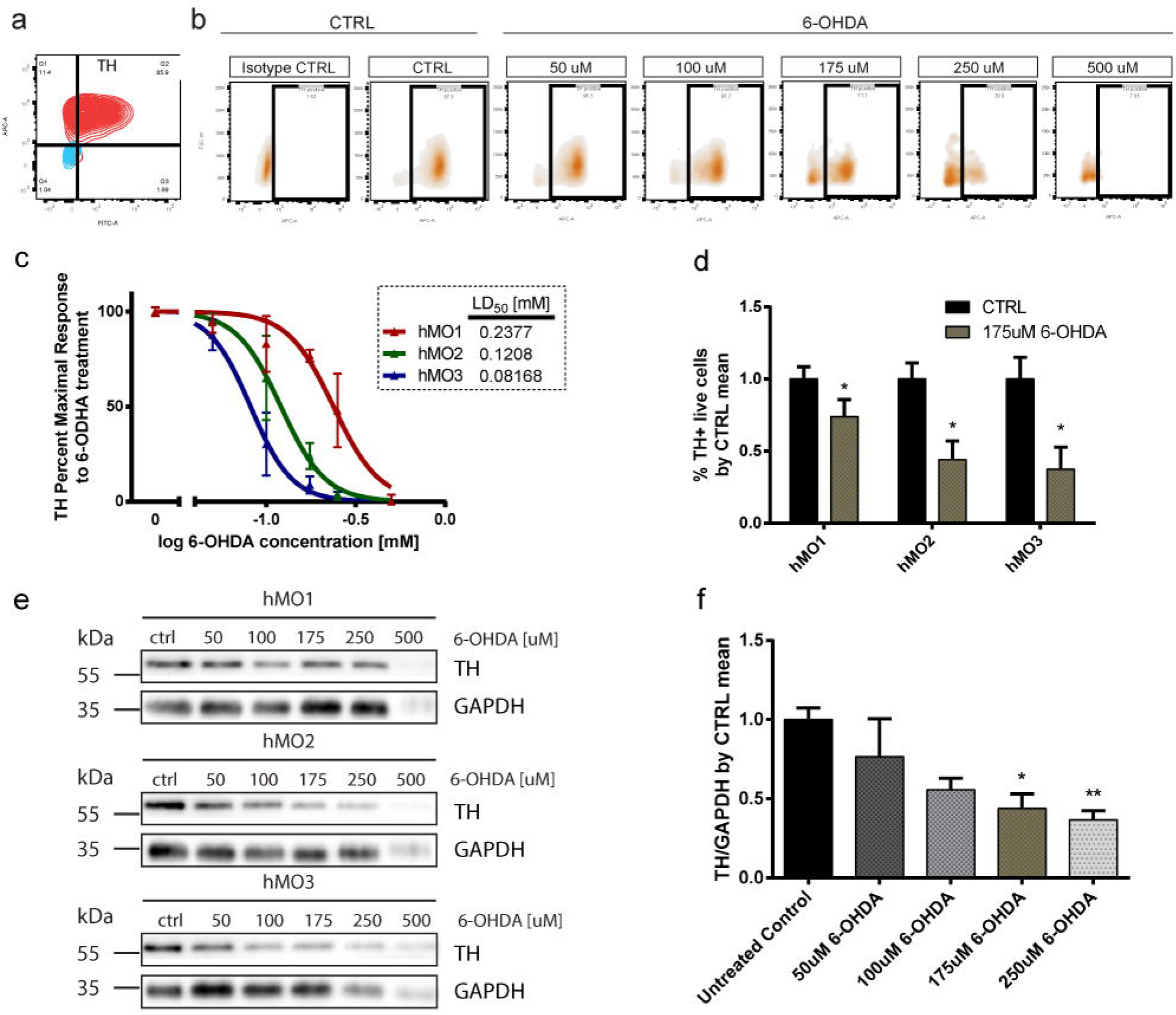
6-OHDA-induced degeneration of dopaminergic neurons. a) Representative flow-cytometry gating setup for TH b) Representative flow-cytometry analysis showing a decrease in TH+ live cells at different concentration of 6-OHDA ranging from 50 to 500µM. c) 6-OHDA dose-response curves fitted to the data of B. d) Barplot showing a robust decrease in the amount of TH+ cells at a 6-OHDA concentration of 175µM. Data obtained for each cell line from seven independent organoid batches and 6-OHDA treatments. Error bars represent mean + SEM. *p< 0.05 e) Western Blot revealing a concentration-dependent decrease of TH protein upon 6-OHDA treatment. f) Quantification of d, normalized to the mean of the untreated controls of nine organoids derived from three independent lines. Error bars represent mean + SEM. *p< 0.05, **p<0.005

### High-content imaging platform

We next examined the effect of 6-OHDA on the neuronal network within midbrain organoids using image-based cell profiling. We developed a high-content imaging platform to automatically acquire, process and analyze images from organoid sections stained for neuron-specific Class III β-tubulin (TUJ1), microtubule-associated protein 2 (MAP2) and dopaminergic neuronal marker TH. We subdivided organoid sections into center (5-6 80µm sections of the organoid core) and border (non-center) sections (Fig. 3a, Fig. S2a) prior to immunostaining. This step is essential to correct for spatial asymmetry in architecturally complex organoids, with a dense core and neurites reaching out in the periphery. We acquired 12-16 area scans and 30 planes per organoid section using an automated confocal microscope (Fig. 3b). The acquired images were stitched in MATLAB (Fig. 3c) and the amount of neurons and dopaminergic neurons was quantified by normalizing to Hoechst positive nuclei. As expected, in untreated organoids we saw a significant difference between border and center sections after normalization (Fig. S2b) due to the nuclear density in the center of the organoid (Fig. S2c). Hence, we analyzed border and center sections separately or corrected for the variation of the section by normalization.

**Fig. 3:**
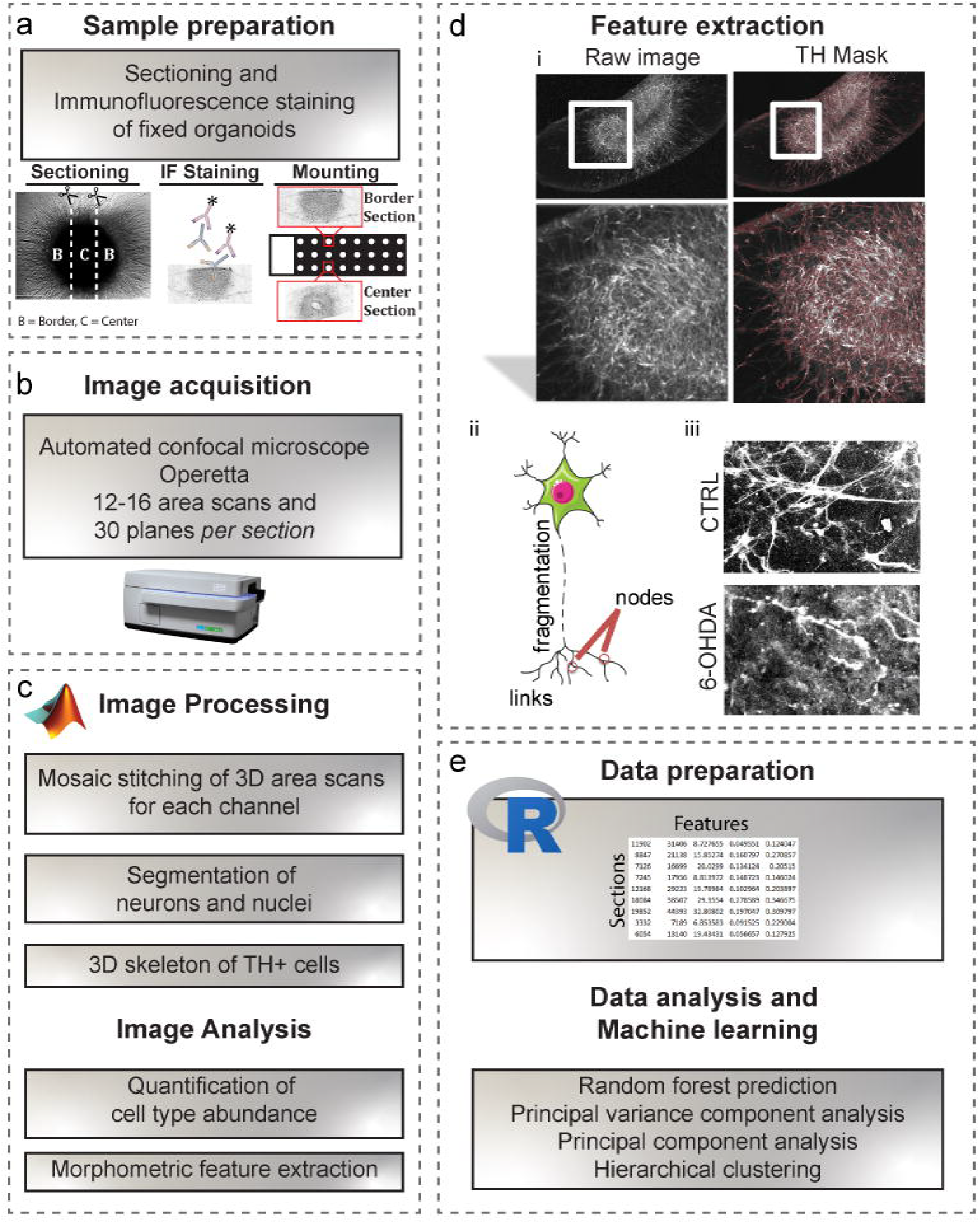
High-Content Image Analysis Workflow. a) Sample preparation. Organoids were sectioned and prior to immnuofluorescence staining separated into border and center sections. Organoid sections were mounted on an object slide containing a grid for automated image acquisition. b) Image acquisition. 12-16 area scans in 30 planes were acquired using an automated confocal microscope. c) Images were exported in MATLAB and area scans were stitched. On the obtained 3D image, masks were generated for dopaminergic neurons in order to quantify cell type abundance and morphometric features. d) Feature extraction. Dopaminergic neuronal complexity was quantified by extracting cellular features such as neurite nodes, links, and fragmentation. i) Mask generation, ii) Schematic view of the extracted features, iii) Example of neurite fragmentation and decreased cellular complexity after 6-OHDA treatment. e) High-content data analysis in R. The obtained data from MATLAB was exported into R and further processed for machine learning-based prediciton of neurotoxicity and neuroprotection.

### Dopaminergic neurons within midbrain organoids show typical signs of degeneration

Upon 6-OHDA treatment, the overall amount of neurons, positive for the neuronal markers TUJ1 and MAP2 remained unaltered (Fig. 4a, b). On the contrary, the amount of TH+ dopaminergic neurons decreased significantly (Fig. 4c). We next computed a 3D mask for TH+ cells using edge-detection methods of image processing. Further, we generated a 3D skeleton of the dopaminergic neuronal network in order to extract features such as nodes (dendritic and axonal points of branching) and links (total number of branches), as well as neurite fragmentation using erosion operations (Fig. 3c, d, Table S2). 6-OHDA treatment leads to a significant decrease in the complexity of dopaminergic neurons and an increase in the amount of fragmented neurites (Fig. 4d-f, Fig. S1c, Fig. S3-5).

**Fig. 4:**
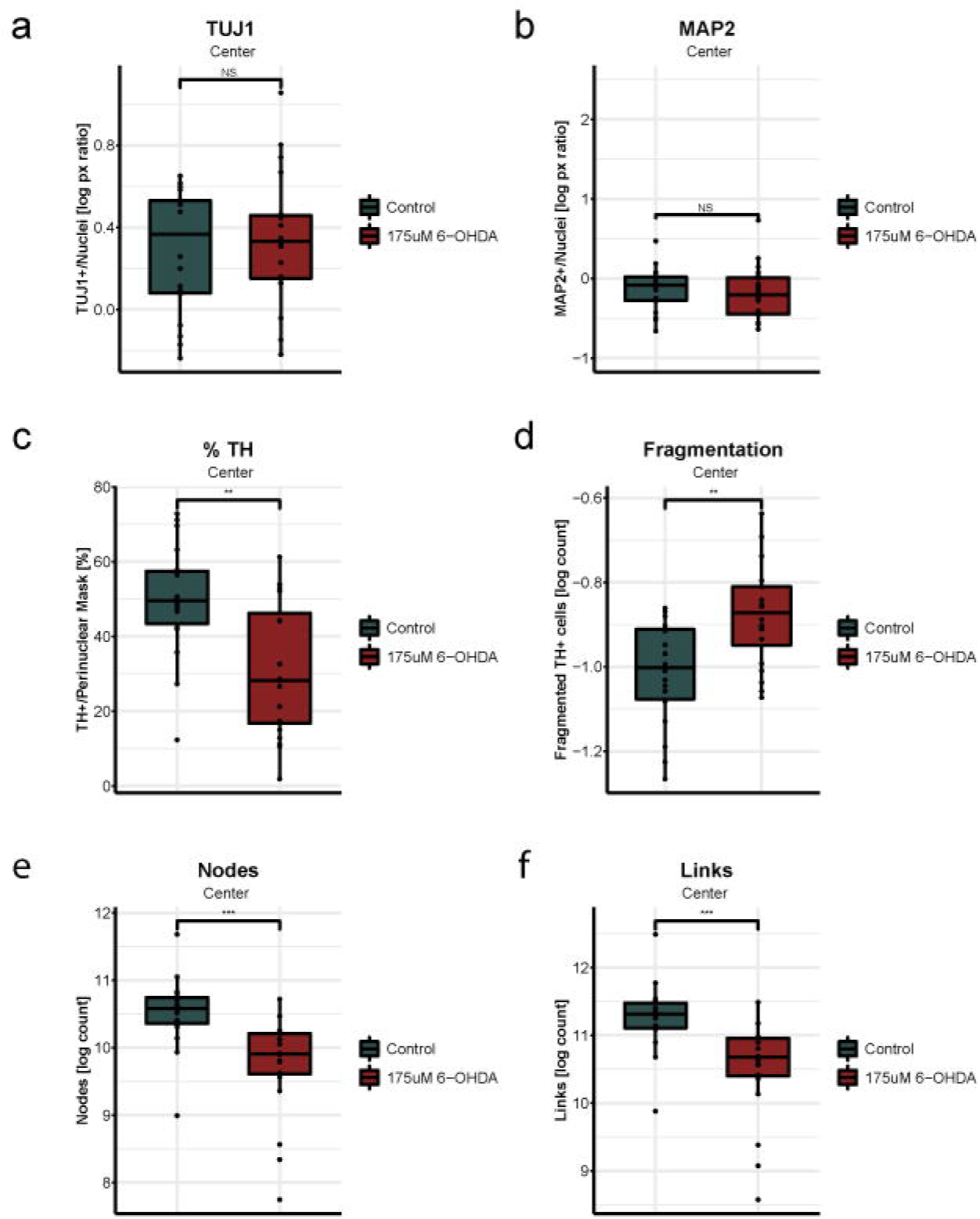
Specific degeneration of dopaminergic neurons after 6-OHDA treatment. a) The overall amount of TUJ1+ neurons is unaffected from the treatment. b) The overall amount of MAP2+ neurons is unaffected from the treatment. c) Dopaminergic neurons degenerate after 6-OHDA treatment. %TH: Total count of TH+ cells. d-f) 6-OHDA treatment leads to impaired neuronal complexity as indicated by increased neurite fragmentation, decreased number of nodes (branch origin and end-point) and increased numbers of links (neurite branches). Data obtained from four independent organoid batches and 6-OHDA treatments from three cell lines. Wilcoxon rank sum test, *p<0.05, **p<0.01, ***p<0.001

### Random forest prediction of neurotoxicity

We next used a ML approach to build a classifier able to discriminate between CTRL and 6-OHDA-treated organoids; and consequently identify the measurements that describe the largest difference between the two conditions. We trained a random forest (RF) algorithm with a ten-time 5-fold cross-validation procedure in order to ensure an unbiased estimation of the model performance. We first applied our strategy to the raw/unprocessed data. The generated model achieved on average a classification accuracy of 75%. The prediction was mainly influenced by dopaminergic features, with the amount of TH+ cells / live cells, TH skeleton, and TH fragmentation as most important measurements (Table 1, Figure S6a_i_,-b_i_).

**Table 1:** Feature-based classification results before and after data normalization

| **Classification**<br>**Algorithm** | **Data**<br>**processing** | **Normalized to** | **Accuracy**<br>**(%)** | **Sensitivity**<br>**(%)** | **Specificity**<br>**(%)** |
| --- | --- | --- | --- | --- | --- |
| RF | Raw | - | 75 | $77 \pm 3$ | $72 \pm 2$ |
| RF | Z-score | E, C, S | 86 | $87 \pm 3$ | $85 \pm 3$ |

Knowing that prediction power of ML models highly depends on data quality, we attempted to remove highly variable and for the biological effect irrelevant experimental factors (organoid batch, cell line and section). We first assessed the contribution of those factors along with the treatment (sample condition) to the variability observed in the data. We used a principal variance component analysis (PVCA) (Bushel, 2013) and observed significant contribution of each factor (Fig. S6c_i_). Building on this, we investigated whether we could improve classification accuracy by normalizing the data. We performed a z-score transformation across the entire dataset for (within each combination of) experiment (4 independent organoid batches and treatments), cell line (hMO1, hMO2, hMO3) and section (border, center). Normalization strongly improved the classification accuracy of the RF model to 86%, while lowering the variance described by experimental conditions (Table 1, Fig. S6a_ii_-c_ii_). Consistent with this, we observed a clear separation between control and 6-OHDA treatment using hierarchical cluster analysis (Fig. 5a), as well as principal component analysis (Fig. 5b). This result suggests that by optimizing data processing strategies, we can robustly predict neurotoxicity using a ML approach on complex high-content image analysis data from human brain organoids.

**Fig. 5:**
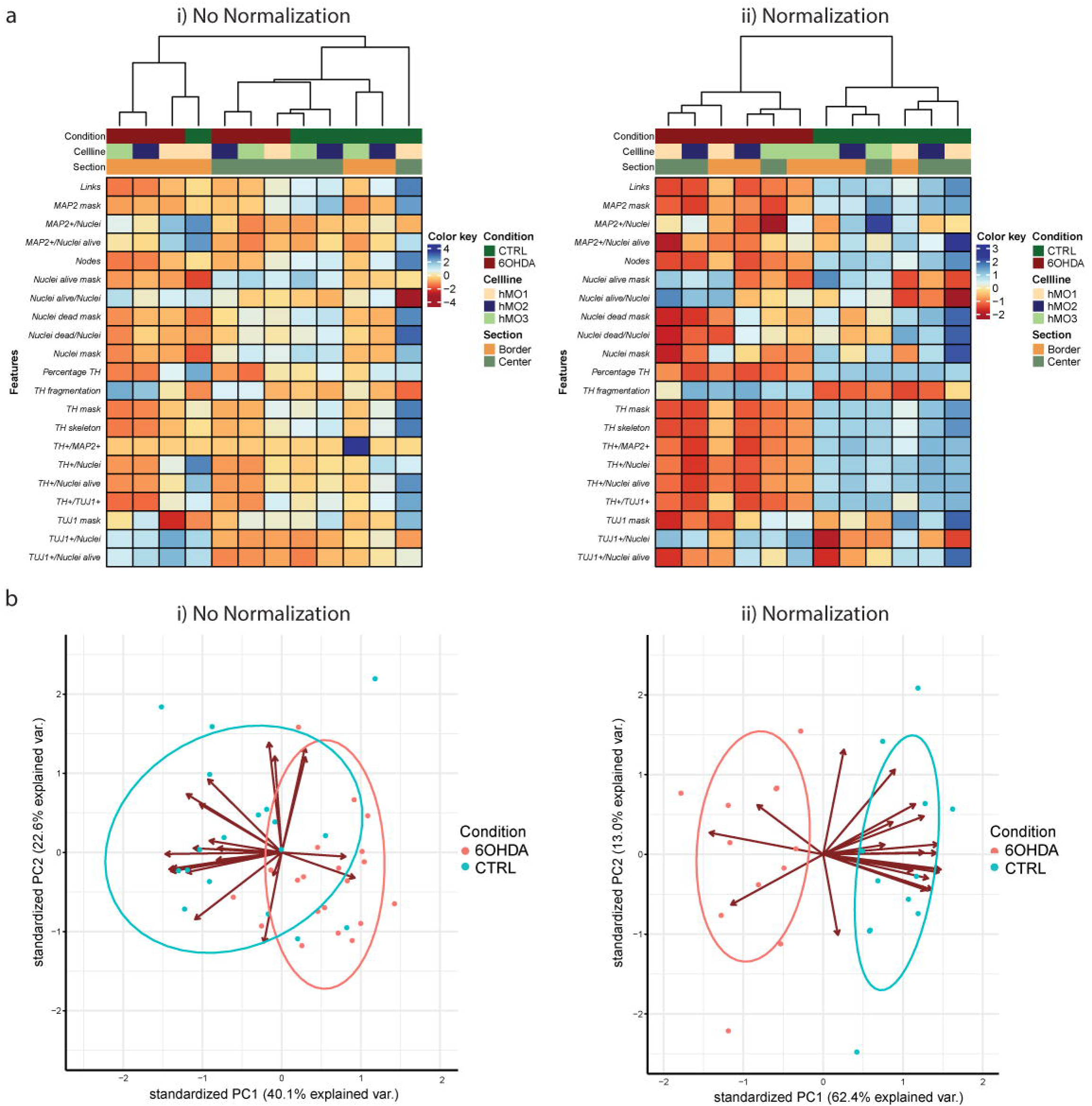
Hierarchical cluster analysis and dimensionality reduction. a) Hierarchical clustering using Ward’s minimum variance method without (i) and with data normalization (ii). b) Principal component analysis before (i) and after data normalization (ii).

## Discussion

In recent years, advanced 3D *in vitro* brain models, termed brain organoids, have been extensively developed to study neurological disorders (Bershteyn et al., 2017; Lancaster et al., 2013; Li et al., 2017; Mariani et al., 2015; Qian et al., 2016), suggesting that brain organoids could also be relevant for modeling of neurodegenerative diseases. 3D culture conditions have been shown to provide the complex environment necessary for extracellular protein aggregation to model Aβ and Tau pathologies (Choi et al., 2014; Lee et al., 2016). The development of regionally restricted midbrain-specific organoids suggests their potential to model Parkinson’s disease (Jo et al., 2016; Monzel et al., 2017). Two recent studies show that genetically modified and patient-derived midbrain organoids harboring the PD-associated LRRK2-G2019S mutation manifest degenerative phenotypes and decreased complexity of dopaminergic neurons (Kim et al., 2019; Smits et al., 2019).

In this study, we have used the catecholaminergic neurotoxin 6-OHDA to target specifically the dopaminergic system. Due to its structural similarity with endogenous dopamine, 6-OHDA enters the dopaminergic neuron via the dopamine transporter, leading to an accumulation of the toxin in the neuron. Because of its selectivity for dopaminergic neurons, 6-OHDA is the drug most frequently used to induce neurodegeneration of the nigrostriatal system in animal models. To date, three mechanisms of action have been proposed for the neurotoxic effect of 6-OHDA. 1) Auto-oxidation of 6-OHDA generating cytotoxic H_2_O_2_, reactive oxygen species (ROS) and catecholamine quinones, causing severe oxidative stress, 2) enzymatic conversion of 6-OHDA to hydrogen peroxide via monoamine oxidase (Simola et al., 2007), and 3) direct inhibition of mitochondrial respiratory chain complex I (Glinka and Youdim, 1995). The resulting oxidative stress is followed by the alteration of cellular homeostasis and neuronal damage, leading to cell death. 6-OHDA has been suggested as a putative neurotoxic environmental factor in the pathogenesis of PD (Jellinger et al., 1995), based on the occurrence of 6-OHDA in human brain (Curtius et al., 1974), as well as in urine of PD patients (Andrew et al., 1993).

To assess neuronal damage in the human midbrain organoid system, among other methods we used microscopy-based phenotyping. However, compared to 2D monolayer cultures, organoids exhibit an architecturally complex heterotypic spatial organization. Typically, multiple cell types like astrocytes, oligodendrocytes, stem cells and neurons are arranged in close proximity in the 3D space, the latter one expanding long neurites in the surrogate matrix. This complexity makes it utterly difficult to measure neuronal complexity, such as branching and thickness of neurites and to quantify these measurements. Hence, the use of powerful image processing algorithms is necessary to extract morphometric features accurately on the single cell level. However, high-content microscopy generates large amounts of multi-parametric data. The technological advances of high imaging throughput, precise analytical frameworks with high-performance computation opens new avenues for phenotypic profiling on the single-cell level in brain organoids. In combination with a powerful machine learning approach for the analysis of multivariate profiling data, we were able to predict neurotoxin-induced perturbations in the human midbrain organoid system. Random forest by design is a well-established technique for reducing predictive variability, preventing overfitting and achieving high classification accuracy (Parmar et al., 2019). Importantly, random forest gives estimates of which variables are most important in the classification (Breiman, 2001). Moreover, using PVCA, we were able to identify the contribution of experimental factors to the total variance and design optimized data normalization approaches to improve predictability. This supports the concept of using image-based profiling studies in organoid models to identify drugs that modulate phenotypes. We suggest that organoids have the potential to be used as a platform from target identification to toxicity prediction using machine learning-assisted high-content image-based cell profiling.

## Experimental Procedures

### Cell Culture

Human NESC lines from three female healthy individuals were derived as described in (Reinhardt et al., 2013) from human iPSCs (Table S1). Human NESCs were cultured on Matrigel-coated plates in N2B27 media supplemented with 3 µM CHIR-99021 (Axon Medchem), 0.75 µM purmorphamine (Enzo Life Science) and 150 µM ascorbic acid (Sigma) (referred to as N2B27 maintenance media) as previously described (Reinhardt et al., 2013). N2B27 medium consists of DMEM-F12 (Invitrogen)/Neurobasal (Invitrogen) 50:50 with 1:200 N2 supplement (Invitrogen), 1:100 B27 supplement lacking Vitamin A (Invitrogen), 1 % L-glutamine and 1 % penicillin/streptomycin (Invitrogen). Midbrain organoids were generated with 9000 cells exactly as described previously (Monzel et al., 2017) with the exeption that Geltrex was used instead of Matrigel as extracellular matrix. hNESCs were plated from single cell suspension following accutase treatment and cultured for 6 days in ultralow attachment 96 well plates (CORNING) in N2B27 maintenance media. At day 8, the NESC spheroids were embedded into droplets of Geltrex, and cultured in non-treated tissue culture 24 well plates (CELTREAT). One organoid was kept per well. At day 10, the differentiation into midbrain dopaminergic neurons was initiated with N2B27 media supplemented with 10 ng/ml hBDNF (Peprotech), 10 ng/ml hGDNF (Peprotech), 500 µM dbcAMP (Peprotech), 200 µM ascorbic acid (Sigma), and 1 ng/ml TGF-β3 (Peprotech). 1 µM purmorphamine (Enzo Life Science) was added to this medium for an additional 6 days. At day 14, organoids were place on an orbital shaker (IKA) rotating at 80 rpm, and media was changed every 3 to 4 days until the treatment.

### Cytotoxicity

In order to identify, which 6-OHDA concentration leads to a significant reduction in the amount of dopaminergic neurons, organoids were treated with 50 μM, 100 μM, 175 μM, 250 μM and 500 μM 6OHDA (Sigma) after five weeks of organoid culture. Organoids were cultured in N2 Medium (DMEM/F12 with 1% N2 supplement and 1% penicillin/streptomycin/glutamine), supplemented with 1:500 DMSO only, or with 6-OHDA with 1:500 DMSO. DMSO was always added to the medium in case a compound treatment requires the use of DMSO, which in our experiments diminished the effect of 6-OHDA due to its antioxidant properties. Due to the instability and rapid auto-oxidation of 6-OHDA, we performed a double-treatment on two sequential days. Two days later, medium was changed against N2B27 differentiation and organoids were kept under usual culture conditions on an orbital shaker. Six days later, organoids were fixed (IF staining), frozen at −80°C (Western blot) or dissociated (Flow cyotmetry). During the treatment, the matrix around the organoids blackened due to the spontaneous oxidation of the compound.

### Flow cytometry

For flow cytometry analysis, six organoids for each cell line and treatment condition were dissociated to single cells by incubation in 0.18% Papain (Sigma), 0.04% EDTA (Sigma) and 0.04% L-Cystein (Sigma) dissolved in DMEM-F12 (Invitrogen) at 37 °C under dynamic conditions until the Geltrex was completely removed (2-3 h). Afterwards, organoids were treated with Accutase (Sigma) for another 1-2 h at 37 °C under dynamic conditions. During Accutase treatment, organoids were carefully dissociated by pipetting, first with a 1000 μl pipette and then with a 200 μl pipette. Prior to fixation, the single cells were washed in cold 1x PBS and stained for live/dead cells using eBioscience™ Fixable Viability Dye eFluor™ 450 for 30 min at 4 °C on a rotor. For intracellular staining, transcription factor buffer set (BD Bioscience) was used according to the manufacturer’s instructions. After fixation, cells were filtered through a 5 ml polystyrene round-bottom tube with cell-strainer cap (Corning). 100,000 cells were used per sample and antibodies were used at the following dilutions: chicken anti-TH (1:50, Abcam) and rabbit anti-TUJ1 (1:500, Covance). Threshold gates were set with the same samples that were stained with the following isotype control antibodies: Normal chicken IgY Control (R&D Systems), and normal rabbit IgG control (Santa Cruz Biotechnology). Isotype control antibodies were used at the same concentration as the detection antibody. All secondary antibodies (Invitrogen) were conjugated to Alexa Fluor fluorochromes. Flow cytometry was performed by using BD LSRFortessa Cell Analyzer and data were analyzed and represented by FlowJo software.

### Western Blot

For Western Blot, one organoid per condition was lysed with Urea Buffer (7M Urea, 2M Thiourea, 2% CHAPS, 1% DTT (w/v) and Complete protease inhibitor cocktail (Roche) in MilliQ water). The cell lysates were centrifuged for 10 min at the highest speed at 4°C. The supernatant was mixed with sample buffer and boiled at 95°C for 5 min. The protein concentration of the boiled samples was determined using a protein quantification assay kit (Macherey Nagel). After adjustment to equal protein concentrations, the samples were subjected to sodium dodecyl sulphate-polyacrylamide gel electrophoresis (SDS-PAGE) and western blotting. The lysates were size-separated by electrophoresis and transferred to nitrocellulose membranes using iBlot™ 2 Gel Transfer Device (Thermo Fischer). The equal loading of the blotted protein was verified by Ponceau S (Sigma) staining. Subsequently, membranes were blocked for 1 h at RT in 5% skimmed milk powder and 0.2% Tween in PBS before incubating overnight at 4◦C with the primary antibodies mouse anti-TH (1:1000, Millipore) and rabbit anti-GAPDH (1:1000, Abcam). Horseradish peroxidase conjugated secondary antibodies and enhanced chemiluminescence reagents (ECL kit) (GE Healthcare) were used for detection. Western blots were analyzed using ImageJ software.

### Immunofluorescence stainings

At day 42, organoids were fixed with 4 % paraformaldehyde overnight at RT and washed 3x with PBS for 15 min. Two organoids per cell line, condition and experiment were embedded in 3% low-melting point agarose in PBS and incubated for 20 min at 40 °C, followed by 30 min incubation at RT. 80 µm sections were cut using a vibratome (Leica VT1000s), and sections were separated into “border” and “center” sections. The sections were permeabilized and blocked with 0.5 % Triton X-100, 2.5 % normal goat or donkey serum, 2.5 % BSA, and 0.1 % sodium azide. Sections were incubated on a shaker for 48 h at 4 °C with primary antibodies in the blocking buffer containing 0.1% Triton X-100 at the following dilutions: rabbit anti-TH (1:1000, Abcam), chicken anti TUJ1 (1:1000, Millipore) and mouse anti-MAP2 (1:200, Millipore). After incubation with the primary antibodies, sections were washed three times in 0.01 % Triton X-100 and incubated with the secondary antibodies (1:1000) including a Hoechst 33342 counterstaining for nuclei in blocking buffer with 0.01 % Triton X-100. All secondary antibodies (Invitrogen) were conjugated to Alexa Fluor fluorochromes. Fluorescence images were acquired on Operetta confocal microscope (Perkin Elmer) with a 20x Objective (16-20 area scans, 25 z-planes).

### Image processing and analysis

Immunofluorescence 3D images of each organoid section in four channels were processed and analyzed in Matlab (2017a, Mathworks) using a custom image-analysis algorithm.

First, mosaic stitching was performed for each channel by computing normalized cross correlations between overlapping image sections. Positions of the local maxima were used to return x and y offsets for the positioning of image tiles in the stitched mosaic image.

In the stitched image, nuclei were segmented via difference of Gaussians. In brief, a foreground image was computed by convolving the raw Hoechst channel with a Gaussian filter of size 21 and standard deviation 1. The background image defined as the raw Hoechst channel convolved with a Gaussian filter of size 21 and standard deviation 3 was subtracted from the foreground image. The nuclei mask was defined by pixels with gray tone values larger than 20. Furthermore, only connected components with at least 20 pixels were kept. For the quantification of dead nuclei, the raw Hoechst channel was convolved with a Gaussian filter of size 11 and standard deviation 1. All pixels with a gray tone value larger than 1800, were considered as pyknotic nuclei pixels, which show an increased fluorescence intensity in Hoechst. Relative quantification of live cells was calculated by subtracting the pyknotic nuclei pixels from the total number of nuclei pixels.

Dopaminergic neurons in the TH channel were segmented using Fourier transform. For high pass frequency filtering, a butterworth filter with cutoff frequency 7 and order 1 was applied to each image plane and the mask was defined by pixels with gray tone values larger than 0.003. Connected components with at least 1000 pixels were kept and the mask was further refined by applying a 3D median filter (3×3×3). Similarly, neurons in the TUJ1 and MAP2 channel were segmented using Fourier transform with cutoff thresholds of 0.0015 and 500 pixels (Marques, 2011).

To extract morphometric features from dopaminergic neurons, a 3D skeleton was generated from the TH mask. The algorithm used for skeletonization is based on homotopic thinning with parallel topology consistency checking (Kerschnitzki et al., 2013; Lee et al., 1994). The resulting skeleton was converted into a network graph describing nodes (branching points) and links (branches). To analyze neuronal fragmentation, the mask in the TH channels was eroded. The underlying theory is that the surface of fragmented objects is larger than the surface of non-fragmented objects as compared to their cumulated volumes. The erosion was performed using a 3D spherical structuring element with a radius of 1 pixel.

Additionally, the amount of TH+ cells was quantified by identifying separated nuclei that have adjacent TH+ pixels, as opposed to a pixel count. First, big interconnected objects were removed from the nucleus mask (> 10000 pixels). This step removes the nuclei from the dense inner core of the organoid, where nuclei cannot be separated. The resulting nuclei were dilated with a disk-shaped structuring element of 4 pixels, and an additional spherical structuring element of size 1. For single cell analysis, an overlap of a dilated nucleus with TH+ pixels was then counted as one TH positive cell.

The extracted features are summarized and described in detail in **Table 2**.

### Data analysis

We assigned additional metadata to the obtained high-content image analysis data. The metadata described the origin of every organoid section (*Cellline* (hMO1, 2, 3), *Experiment* (organoid batch/-treatment), *Replicate* (1, 2), *Condition* (CTRL, 6-OHDA), and *Section* (border/center). The data was imported into R software (R version 3.5.1 --"Feather Spray") and further processed. First, data from *Cellline*, *Replicate*, *Experiment, Condition* and *Section* was grouped and the mean of the measurements was calculated. This step was performed in order to combine data from neighboring sections. We used data-analysis strategies as suggested in (Caicedo et al., 2017).

### Classification model based on RF

We used random forest (Breiman, 2001) to model the treatment condition of the organoids. RF is an ensemble method based on classification and regression tree (CART). It is very popular since derived models usually reach high performances (Touw et al., 2013). RF aims at creating several trees that will be used to predict the class of a new sample. The final decision is made through a voting system. For classification, each tree is built by iteratively selecting a set of features and choose among them the candidate that maximizes the separation between samples of different conditions. This process is repeated until either the separation is perfect or the maximum number of iteration is reached or no improvement is possible. Note that CARTs are non-parametric. Features are discretized according to the value that gives the best sample splitting. CART therefore generates binary tree. To perform RF, we used the randomForest function of randomForest R package (Liaw and Wiener, 2002). To assess the prediction accuracy of RF models, we implemented a standard 5-fold cross-validation procedure. A key advantage of RF resides in its automatic evaluation of the importance of each feature. For a given feature, we estimated its global feature importance by averaging the importance measures obtained from 50 runs of RF models.

## Data Availability

The data that was used to build up the model is openly available at DOI: 10.17881/lcsb.20191309.02.

## Code Availability

The Matlab and R scripts for the image and data analysis are available on github. The R scripts for RF classification are available on gitlab.

## Acknowledgements

We thank Prof. Dr. Hans R. Schöler of the Max Planck Institute, Prof. Dr. Thomas Gasser of the Hertie Institute in Tübingen and the Coriell Institute for providing cell lines. Microscopy was supported by the LCSB bio-imaging platform. We thank the Disease Modeling Screening Platform from LCSB and LIH for their help with performing automated and high-throughput procedures. This work was supported by a Proof-of-Concept grant from the Fonds National de la Recherche (FNR) Luxembourg (FNR/PoC16/11559169). Further, this is an EU Joint Program - Neurodegenerative Disease Research (JPND) project (INTER/JPND/14/02; INTER/JPND/15/11092422) receiving funding from the FNR. A.S.M. and I.R. were supported by FNR Aides à la Formation-Recherche (AFR). A.S.M. received support from the Lush prize 2017.

Finally, we also thank the private donors who support our work at the Luxembourg Centre for Systems Biomedicine.

## Author Contributions

A.S.M. designed experiments, prepared the figures, performed image and data analysis and wrote the original draft. K.H. and I.R. designed experiments, and prepared the figures. T.K.M. performed RF classification, prepared figures and wrote the original draft. P.L., S.L.N., and A.Z. designed experiments. P.A. and S.B. designed image analysis, and edited the manuscript. R.K., F.A., J.C.S. conceived and supervised the project, designed the experiments, and edited the manuscript.

## References

Andrew, R., Watson, D.G., Best, S.A., Midgley, J.M., Wenlong, H., and Petty, R.K. (1993). The determination of hydroxydopamines and other trace amines in the urine of parkinsonian patients and normal controls. Neurochem. Res. 18, 1175–1177.

Bellou, V., Belbasis, L., Tzoulaki, I., Evangelou, E., and Ioannidis, J.P.A. (2016). Environmental risk factors and Parkinson’s disease: An umbrella review of meta-analyses. Parkinsonism Relat. Disord. 23, 1–9.

Bolognin, S., Fossépré, M., Qing, X., Jarazo, J., Ščančar, J., Moreno, E.L., Nickels, S.L., Wasner, K., Ouzren, N., Walter, J., et al. (2019). 3D Cultures of Parkinson’s Disease-Specific Dopaminergic Neurons for High Content Phenotyping and Drug Testing. Adv. Sci. 6, 1800927.

Breiman, L. (2001). Random Forests. Mach. Learn. 45, 5–32.

Bushel, P. (2013). Principal Variance Component Analysis (PVCA). Version 1.22.0.

Caicedo, J.C., Cooper, S., Heigwer, F., Warchal, S., Qiu, P., Molnar, C., Vasilevich, A.S., Barry, J.D., Bansal, H.S., Kraus, O., et al. (2017). Data-analysis strategies for image-based cell profiling. Nat. Methods 14, 849–863.

Curtius, H.C., Wolfensberger, M., Steinmann, B., Redweik, U., and Siegfried, J. (1974). Mass fragmentography of dopamine and 6-hydroxydopamine. Application to the determination of dopamine in human brain biopsies from the caudate nucleus. J. Chromatogr. 99, 529–540.

Dorsey, E.R., and Bloem, B.R. (2018). The Parkinson Pandemic—A Call to Action. JAMA Neurol. 75, 9.

Emborg, M.E. (2007). Nonhuman primate models of Parkinson’s disease. Ilar J 48, 339–355.

GBD (2017). Global, regional, and national burden of neurological disorders during 1990-2015: a systematic analysis for the Global Burden of Disease Study 2015. Lancet Neurol. 16, 877–897.

Glinka, Y.Y., and Youdim, M.B.H. (1995). Inhibition of mitochondrial complexes I and IV by 6-hydroxydopamine. Eur. J. Pharmacol. Environ. Toxicol. Pharmacol. 292, 329–332.

Hodge, R.D., Bakken, T.E., Miller, J.A., Smith, K.A., Barkan, E.R., Graybuck, L.T., Close, J.L., Long, B., Johansen, N., Penn, O., et al. (2019). Conserved cell types with divergent features in human versus mouse cortex. Nature 573, 61–68.

Jellinger, K., Linert, L., Kienzl, E., Herlinger, E., and Youdim, M.B. (1995). Chemical evidence for 6-hydroxydopamine to be an endogenous toxic factor in the pathogenesis of Parkinson’s disease. J. Neural Transm. Suppl. 46, 297–314.

Jonsson, G., and Sachs, C. (1975). Actions of 6-hydroxydopamine quinones on catecholamine neurons. J Neurochem 25, 509–516.

Kerschnitzki, M., Kollmannsberger, P., Burghammer, M., Duda, G.N., Weinkamer, R., Wagermaier, W., and Fratzl, P. (2013). Architecture of the osteocyte network correlates with bone material quality. J. Bone Miner. Res. 28, 1837–1845.

Klein, C., and Westenberger, A. (2012). Genetics of Parkinson’s Disease. Cold Spring Harb. Perspect. Med. 2, a008888.

Lee, T.C., Kashyap, R.L., and Chu, C.N. (1994). Building Skeleton Models via 3-D Medial Surface Axis Thinning Algorithms. CVGIP Graph. Model. Image Process. 56, 462–478.

Liaw, A., and Wiener, M. (2002). Classification and Regression by random Forest. R News Vol. 2/3, pp 18–22.

Marques, O. (2011). Practical image and video processing using MATLAB (Wiley-IEEE Press).

Monzel, A.S., Smits, L.M., Hemmer, K., Hachi, S., Moreno, E.L., van Wuellen, T., Jarazo, J., Walter, J., Brüggemann, I., Boussaad, I., et al. (2017). Derivation of Human Midbrain-Specific Organoids from Neuroepithelial Stem Cells. Stem Cell Reports 8, 1144–1154.

Parmar, A., Katariya, R., and Patel, V. (2019). A Review on Random Forest: An Ensemble Classifier. (Springer, Cham), pp. 758–763.

Reinhardt, P., Glatza, M., Hemmer, K., Tsytsyura, Y., Thiel, C.S., Höing, S., Moritz, S., Parga, J.A., Wagner, L., Bruder, J.M., et al. (2013). Derivation and Expansion Using Only Small Molecules of Human Neural Progenitors for Neurodegenerative Disease Modeling. PLoS One.

Scheeder, C., Heigwer, F., and Boutros, M. (2018). Machine learning and image-based profiling in drug discovery. Curr. Opin. Syst. Biol. 10, 43–52.

Schwamborn, J.C. (2018). Is Parkinson’s Disease a Neurodevelopmental Disorder and Will Brain Organoids Help Us to Understand It? Stem Cells Dev. 27, 968–975.

Simola, N., Morelli, M., and Carta, A.R. (2007). The 6-Hydroxydopamine model of parkinson’s disease. Neurotox. Res. 11, 151–167.

Smits, L.M., Reinhardt, L., Reinhardt, P., Glatza, M., Monzel, A.S., Stanslowsky, N., Rosato-Siri, M.D., Zanon, A., Antony, P.M., Bellmann, J., et al. (2019). Modeling Parkinson’s disease in midbrain-like organoids. Npj Park. Dis. 5, 5.

Sulzer, D. (2007). Multiple hit hypotheses for dopamine neuron loss in Parkinson’s disease. Trends Neurosci. 30, 244–250.

Thoenen, H., and Tranzer, J.P. (1968). Chemical sympathectomy by selective destruction of adrenergic nerve endings with 6-Hydroxydopamine. Naunyn Schmiedebergs Arch Exp Pathol Pharmakol 261, 271–288.

Touw, W.G., Bayjanov, J.R., Overmars, L., Backus, L., Boekhorst, J., Wels, M., and van Hijum, S.A.F.T. (2013). Data mining in the Life Sciences with Random Forest: a walk in the park or lost in the jungle? Brief. Bioinform. 14, 315–326.

Ungerstedt, U. (1968). 6-Hydroxy-dopamine induced degeneration of central monoamine neurons. Eur J Pharmacol 5, 107–110.

Wang, H. (2018). Modeling Neurological Diseases With Human Brain Organoids. Front. Synaptic Neurosci. 10, 15.

