## Supplementary Information and Figures for "Machine learning-assisted neurotoxicity prediction in human midbrain organoids"

| Source of hiPSCs | Age at sampling | Gender | hNESC ID | Corresponding human midbrain organoid culture |
|---|---|---|---|---|
| Reinhardt et al. 2013 | 81 | ♀ | 3.0.0.10.0 | hMO1 |
| Reinhardt et al. 2013 | 53 | ♀ | 3.0.0.14.0 | hMO2 |
| Coriell (ND34769) | 68 | ♀ | 3.0.0.24.1 | hMO3 |

**Table S1: Overview of the cell lines used to generate hMOs.**

Human NESCs were derived under 2D conditions from human iPSCs as described in (Reinhardt et al., 2013) and served as a starting population for the generation of hMOs.

| Feature | Description |
|---|---|
| TUJ1 mask | Count of TUJ1 mask pixel |
| TUJ1+/Nuclei | Sum of TUJ1+ pixels / Hoechst |
| TUJ1+/Nuclei alive | Sum of TUJ1+ pixels / nuclear mask of live cells |
| MAP2 mask | Count of MAP2 mask pixel |
| MAP2+/Nuclei | Sum of MAP2+ pixels / Hoechst |
| MAP2+/Nuclei alive | Sum of MAP2+ pixels / nuclear mask of live cells |
| Nuclei mask | Count of nuclear mask pixels |
| Nuclei alive mask | Count of nuclear alive pixels |
| Nuclei dead mask | Count of nuclear dead pixels |
| Nuclei alive/nuclei | Count of nuclear alive pixels / Hoechst |
| Nuclei dead/nuclei | Count of nuclear dead pixels / Hoechst |
| TH mask | Count of TH mask pixel |
| TH+/Nuclei | Sum of TH+ pixels / Hoechst |
| TH+/Nuclei alive | Sum of TH+ pixels / nuclear mask of live cells |
| TH+/TUJ1+ | Sum of TH+ pixels / Sum of TUJ1+ pixels |
| TH+/MAP2+ | Sum of TH+ pixels / Sum of MAP2+ pixels |
| Percentage TH | Percentage of TH+ cells in the perinuclear zone of segmented nuclei |
| TH Skeleton | Count of TH Skeleton pixels |
| TH Nodes | Total number of branchpoints and endpoints in the TH skeleton |
| TH Links | Total number of branches in the TH skeleton |
| TH Fragmentation | Surface to volume ratio of TH mask |

**Table S2: Features from image analysis**

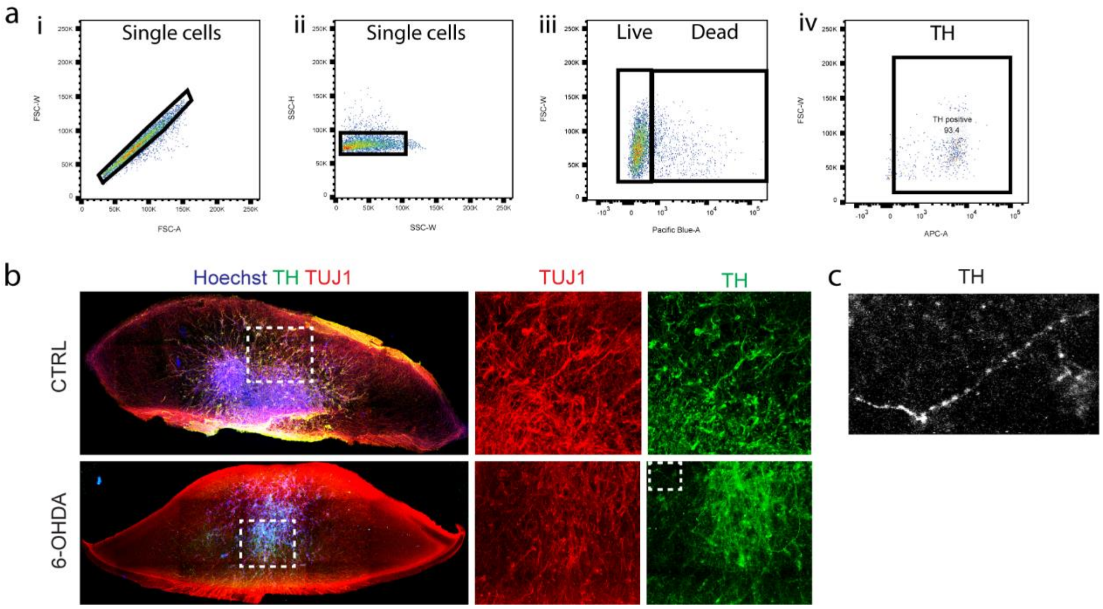


**Fig. S1**

a) Representative flow-cytometry gating setup to single (i,ii), live (iii), and TH+ cells (iv) (related to Fig. 2).

b) Immunofluorescence staining for dopaminergic neuronal marker TH and neuron-specific class III beta Tubulin (TUJ1) in untreated and 6-OHDA treated organoids reveals dopaminergic neurodegeneration upon treatment with 175 μM 6-OHDA.

c) Example of a fragmented TH+ neurite after 6-OHDA treatment.

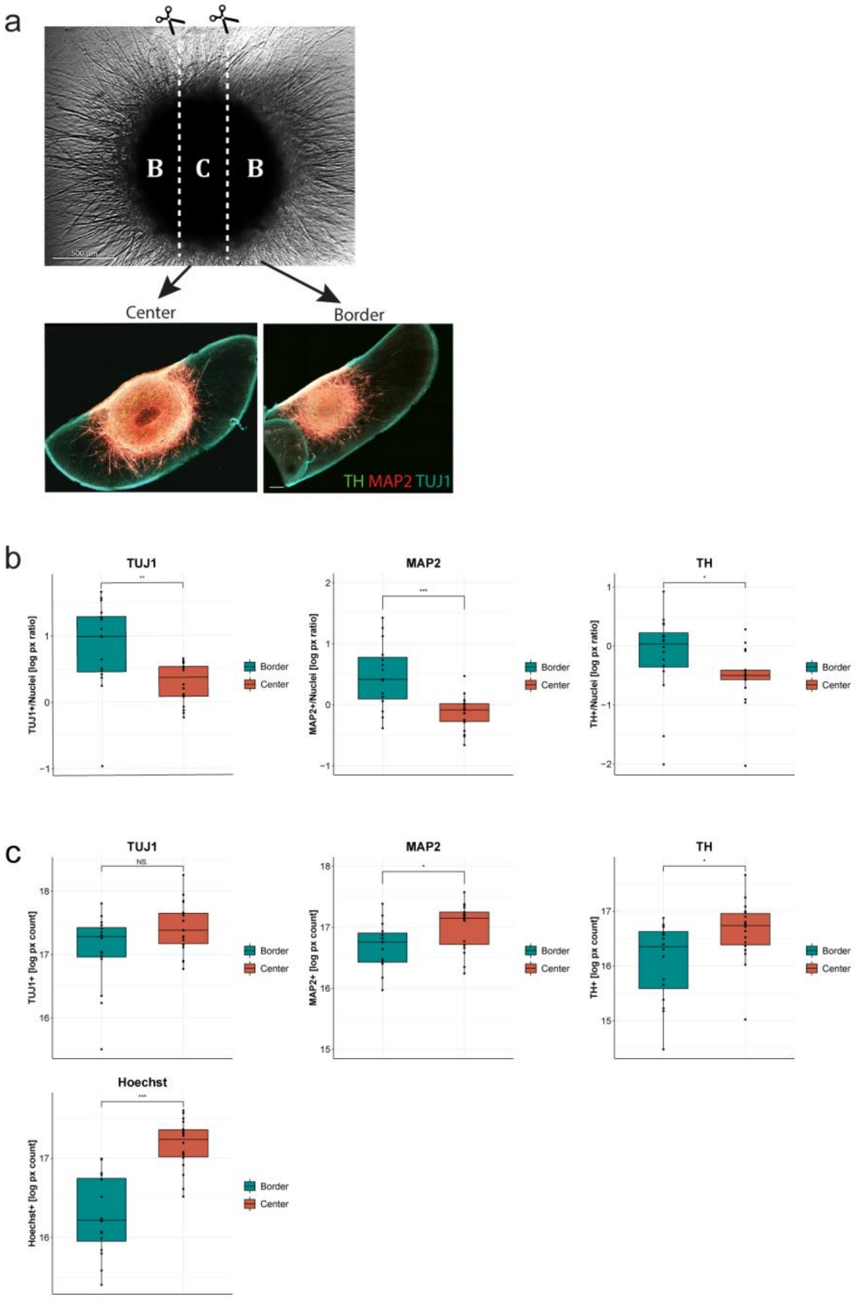


**Fig. S2: Comparison of border and center sections.**

a) Example section for a border and a center region of an organoid stained for TUJ1, MAP2, and TH.

b) Comparison of border (blue) and center (red) sections for TUJ1, MAP2, and TH, normalized to Hoechst.

c) Comparison of border (blue) and center (red) sections for TUJ1, MAP2, and TH pixel count, indicating that the cellular density in center sections is elevated.

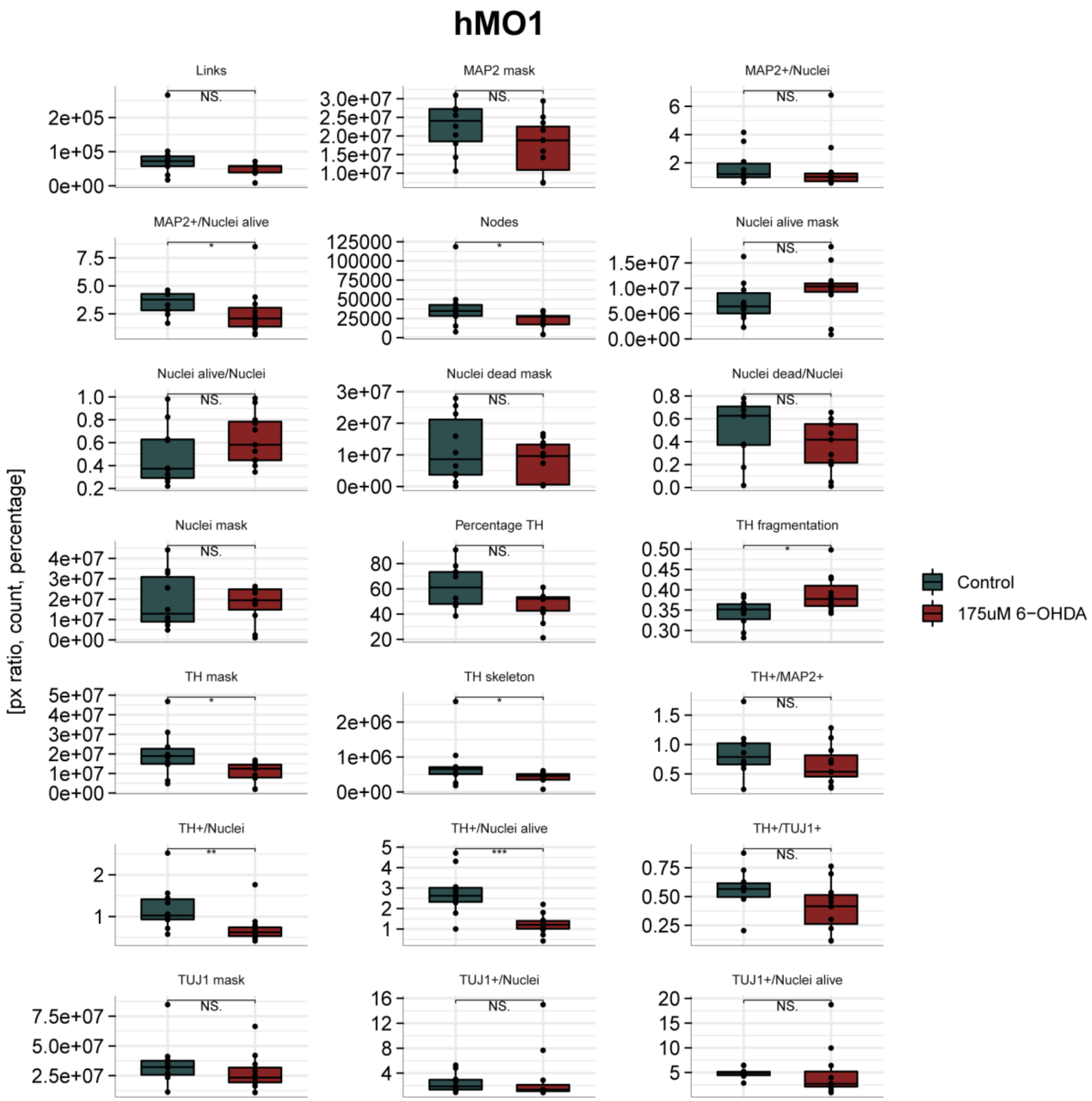


**Fig. S3: Measured morphometric features and cell type abundance of hMO1.**

Raw, unprocessed data of 4 organoid batches and treatments. Border and Center sections pooled. Wilcoxon rank sum test, *p<0.05, **p<0.01, ***p<0.001

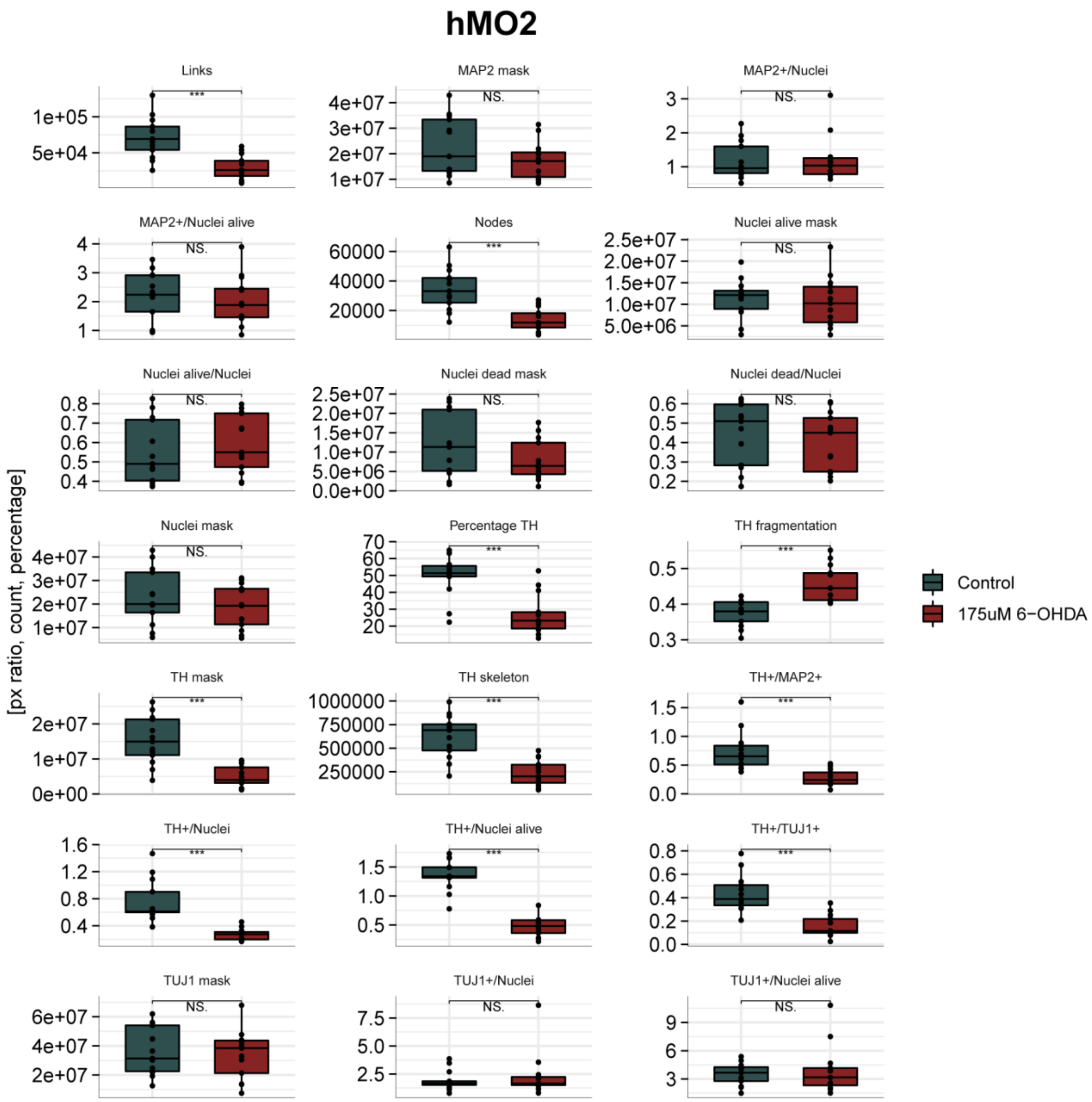


**Fig. S4: Measured morphometric features and cell type abundance of hMO2.**

Raw, unprocessed data of 4 organoid batches and treatments. Border and Center sections pooled. Wilcoxon rank sum test, *p<0.05, **p<0.01, ***p<0.001

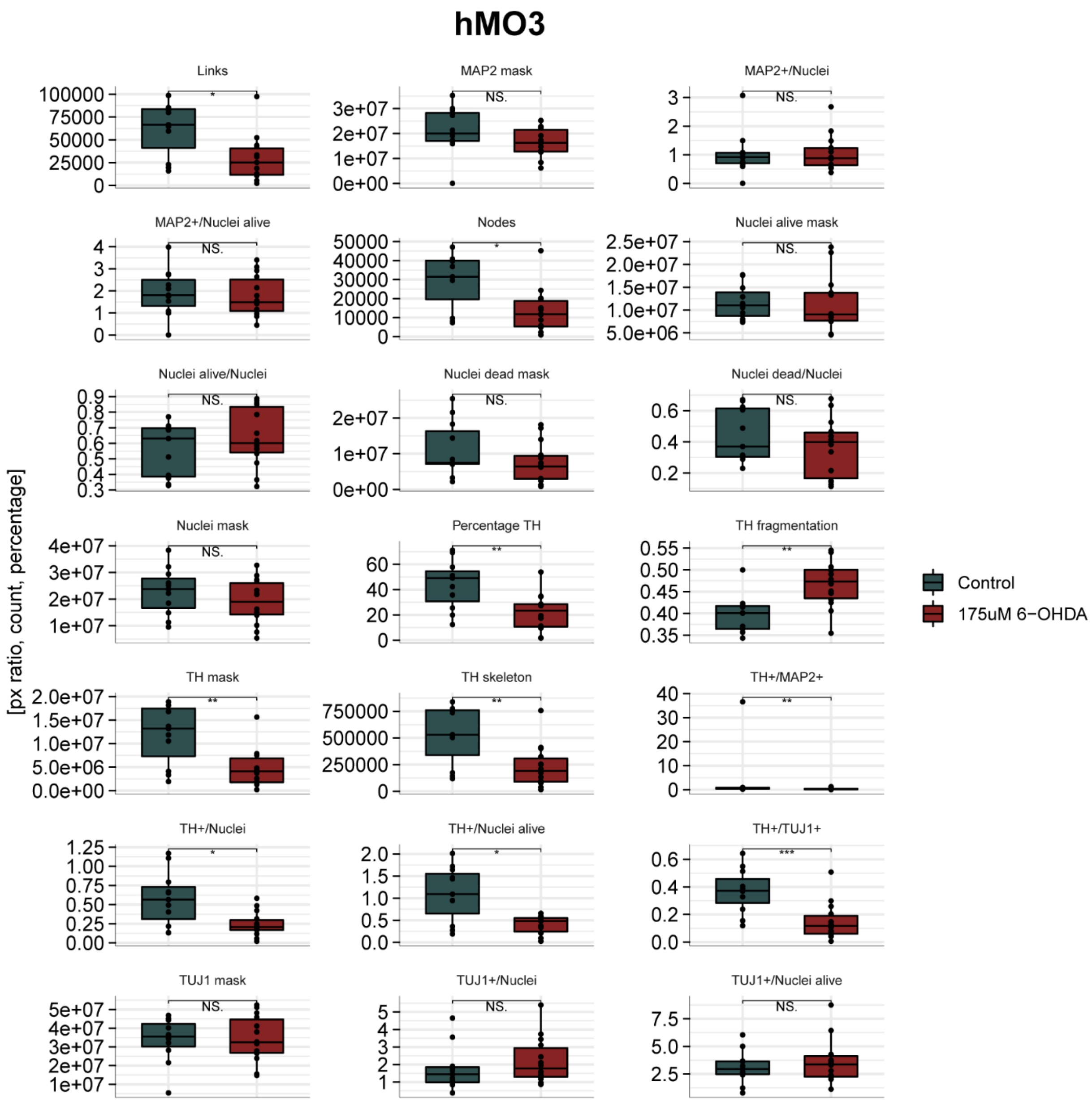


**Fig. S5: Measured morphometric features and cell type abundance of hMO3.**

Raw, unprocessed data of 4 organoid batches and treatments. Border and Center sections pooled. Wilcoxon rank sum test, *p<0.05, **p<0.01, ***p<0.001

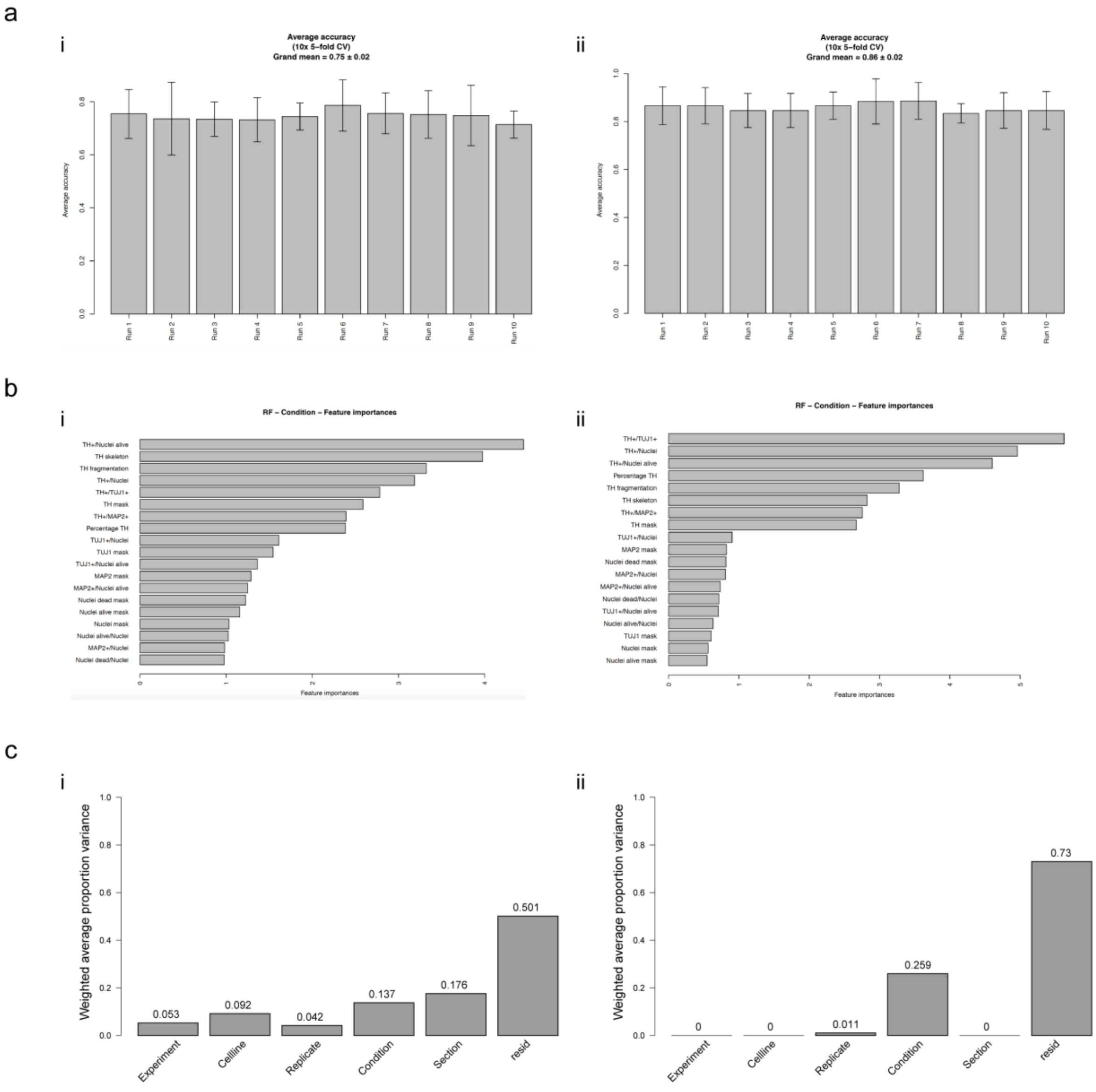


**Fig. S6: Related to Table 1. Comparison between unprocessed (i, left column) and normalized data (ii, right column)**. a) Random forest classification accuracies. Each bar represents the average accuracy over 5-fold cross validation (error bar: ± standard deviation). b) Random forest feature importance reflecting the contribution of each feature to the prediction. c) Principal variance component analysis showing the relative contribution of each experimental factor to the total variance observed in the data.
